# Effects of Overripe Banana Processing Techniques on the Chemical Composition and Attractancy to *Aedes aegypti* and *Aedes albopictus*

**DOI:** 10.64898/2026.11.06.764784

**Authors:** Sara Abdelrahman Abuelmaali, Leow Wen Hui, Sumiyyah Sabar, Nur Amirah Abdul Latif, Wan Rosli Wan Ishak, Ng Yee Vern, Cameron E. Webb, Nurfaeza Abu Kassim

## Abstract

The rising incidence of vector-borne diseases transmitted by mosquitoes has become a significant public health concern. The increasing resistance of mosquitoes to conventional insecticides has prompted the need for alternative control strategies such as Attractive Targeted Sugar Baits (ATSBs). While fruit-based attractants have shown promise, the impact of different production methods on their attractiveness remains underexplored. This study investigates the effect of three production techniques of overripe banana (Musa acuminata) extracts: liquid extraction, syrup concentration, and freeze-drying, on the chemical composition and their attractiveness to *Aedes aegypti* and *Aedes albopictus*. Extracts were infused into hydrogel beads and tested in dual-choice olfactometer assays to mosquito vectors *Ae. aegypti* and *Ae. albopictus*. Chemical profiling was conducted through Gas chromatography-mass spectrometry (GC-MS) and enzymatic sugar analysis. Results showed that all banana extracts were significantly more attractive than the 10% sucrose control. Among the extracts, syrup concentrates showed the highest attraction (PI = 0.58 ± 0.17 for *Ae. aegypti*; 0.58 ± 0.14 for *Ae. albopictus*), followed by freeze-dried (PI = 0.52 ± 0.20; 0.54 ± 0.08) and liquid extracts (PI = −0.40 ± 0.04; 0.49 ± 0.18). GC-MS analysis revealed that freeze-dried powder retained the greatest diversity and abundance of volatile organic compounds (VOCs), especially esters like butyl ethanoate and isoamyl acetate, which are known to influence mosquito behaviour. Although syrup concentration resulted in VOCs degradation, it also generated Maillard reaction products, such as sulphur-containing volatiles, which may enhance olfactory cues. Sugar analysis indicated that both syrup and freeze-dried extracts contained higher levels of sucrose (0.35–0.57 g/100g), glucose (21.20–28.70 g/100g), fructose (14.64–17.89 g/100g), and maltose (9.28–15.56 g/100g), aligning with mosquito preferences. Overall, the findings demonstrate that production techniques significantly influence the chemical composition and attractant potential of overripe banana extracts, offering valuable insights for the development of fruit-based mosquito lures in vector surveillance and control strategies.

## Introduction

Numerous pathogenic viruses continue to be mostly transmitted by mosquitoes, and vector-borne diseases are increasingly prevalent globally. *Aedes* mosquitoes are recognized vectors for the transmission of dengue, yellow fever, chikungunya, and Zika viruses [1, 2, 4, 5]. In 2024, the World Health Organization recorded more than 7.6 billion dengue cases in over 90 countries, including over 16,000 severe cases and 3,000 deaths [6]. *Aedes albopictus* and *Aedes aegypti* are the two predominant *Aedes* species in Malaysia [7,8], identifiable by their body patterns. *Ae. aegypti* is distinguished from *Ae. albopictus* by its white longitudinal stripes and crescent-shaped patterns on the scutum [9], and [10] affirm that, *Ae. albopictus* possesses a solitary white stripe centrally located.

Chemical pesticides are extensively utilized due to their efficacy and rapid action [11,12]. Nonetheless, the frequent and excessive application of pesticides has resulted in resistance in several mosquito populations, hence diminishing the effectiveness of traditional control methods [13,14,15,16]. To resolve this issue, Attractive Targeted Sugar Baits (ATSBs) were created, this approach integrates plant sugars, which are naturally ingested by mosquitoes, with oral insecticides [17,18, 19,8]. Plant-derived attractants such as papaya, pineapple, and sweet alyssum have demonstrated significant appeal to mosquitoes [20,21,22] indicate that boric acid and Spinosad are frequently utilized as the dangerous components in ATSBs. Nonetheless, ATSBs possess a limited lifespan due to the rapid evaporation of attractant chemicals [18,23]; [19]. Alginate–gelatin hydrogel beads (AGHBs) have been proposed as carriers that progressively release pesticides and attractants to enhance stability. AGHBs are environmentally benign and biodegradable, and they enhance the efficacy of sugar baits [24,18,19].

Plant-based attractants, being abundant in carbohydrates and volatile organic compounds (VOCs), naturally draw both male and female mosquitoes. Mosquitoes may select their preferred food sources by their senses of taste, olfaction, and vision, research indicates that *Aedes* species finds guava, mango, and plum juices particularly appetizing [25,8]. *Ae. aegypti* may be particularly attracted to Cavendish bananas (*Musa acuminata*) because of their volatile organic compounds (VOCs) [26].

In 2023, Malaysia generated more than 340,000 metric tons of bananas, affirming its status as a notable producer [27]. Nonetheless, they are classified as some of the most wasted fruits due to their swift decay [28]. Thus, overripe bananas effectively function as a sustainable and cost-effective raw material for mosquito attractants.

Numerous research has identified fruit-based attractants, primarily employing rudimentary extraction methods such as crushing or blending fresh fruit [29,2519]. The impact of different processing techniques on sugars and volatile organic compounds (VOCs) that alter mosquito attraction has not been thoroughly investigated. This study examines how processing techniques, including liquid extraction, syrup concentration, and freeze-drying, alter the chemical content and appeal of overripe banana extracts.

The aim of this study is to evaluate the responses of *Aedes aegypti* and *Aedes albopictus* to hydrogel beads infused with differently processed overripe banana extracts. The study also seeks to identify and characterize the volatile organic compounds (VOCs) present in overripe banana products through multiple analytical approaches, as well as to assess the sugar content of these products and determine the effects of processing on their chemical composition.

## Methods

### Rearing of laboratory strain *Aedes* mosquitoes

Eggs of *Aedes aegypti* and *Aedes albopictus* were taken from the VCRU insectarium at the School of Biological Sciences, Universiti Sains Malaysia. The mosquitoes were reared following the method of [30], under conditions of 27 ± 2 °C temperature, 70 ± 10% humidity, and a 12-hour light and 12-hour dark cycle [23]. First, the eggs were placed in trays filled with clean, dechlorinated water so that they could hatch. After hatching, the larvae were fed every two days with larval food from VCRU. The dirty layer on the water surface was removed.

When the larvae became pupae, they were picked up using a plastic pipette and placed inside rearing cages (30 × 30 × 30 cm) in a small clean water container, adult mosquitoes were given 10% sugar solution soaked in cotton wool. Female adult mosquitoes aged 5–7 days were given a blood meal once a week using mice provided by ARASC, USM. They were starved for one day before blood feeding to help them feed better.

For egg laying, a piece of filter paper soaked with dechlorinated water was put inside each cage. After a few days, the filter papers with eggs were taken out, dried, and stored in a dry container for later use.

### Production techniques of overripe banana

#### Preparation of overripe banana

Cavendish bananas (Musa acuminata) at ripeness stage 5 were bought from Lotus supermarket in Sungai Dua, Penang. The bananas were kept at room temperature (25 °C) and were not treated with any ripening chemicals. They were allowed to continue ripening naturally until they became overripe. The ripeness stage was checked using the peel colour chart described by [31]. When the bananas reached the overripe stage, they were peeled and cut into small pieces for further use.

#### Liquid extraction

The water extraction method for overripe bananas described by [32] was used with a slight modification. About 500 g of overripe banana pulp was blended with distilled water at a 1:3 (w/w) ratio using a Pensonic® PB 3303 blender. The blended mixture was then heated to 50 °C for 30 minutes on a magnetic hotplate stirrer (Joanlab® HS550Pro) to help release the banana components into the water.

After heating, the mixture was poured into 50 mL Falcon tubes and centrifuged at 7000 rpm for 10 minutes. The supernatant (liquid layer) was collected and filtered through filter paper to remove banana fibres. Finally, the clear banana extract was transferred into screw cap bottles and stored at 4 °C until it is needed for further tests and experiments (Appendix C).

#### Syrup concentration

The concentrated banana syrup was prepared using the technique described by [32], with a few modifications. First, 500 g of overripe banana pulp was mixed with distilled water in a 1:3 (w/w) ratio. The combined slurry was put into 50 mL Falcon tubes and centrifuged at 7000 rpm for 10 minutes. The liquid (supernatant) was collected and filtered using filter paper to eliminate the banana fibres. The filtered liquid was then concentrated using a double-boiler technique. The syrup was held at a temperature of 70-80 °C until it reached 75-80 °Bx (Brix value), indicating that most of the water had evaporated (Appendix D).

The thick banana syrup was kept in screw-cap bottles at 4 °C for further analysis and testing.

#### Freeze-drying

About 500 g of overripe banana pulp was blended with distilled water at a 1:3 (w/w) ratio. The blended mixture was then poured into 50 mL Falcon tubes and centrifuged at 7000 rpm for 10 minutes. The liquid (supernatant) was collected and filtered through filter paper.

The clear banana liquid was then poured into ice cream style plastic bags (1.5 × 12 inches) and kept at −80 °C for 48 hours to freeze completely. Before freeze drying, the frozen banana blocks were taken out of the bags and broken into smaller pieces to help the drying process. The banana ice pieces were placed in a freeze dryer (Labconco® FreeZone 6L) and dried at a pressure of 23–45 Pa for 96 hours, with the condenser temperature kept at −51 °C the whole time. After drying, the banana powder was collected and stored at −18 °C for later analysis and experiments (Appendix E).

#### Mosquito attraction bioassay Olfactometer design

This study employed a modified Y-tube olfactometer based on [33] design. The olfactometer consisted of three primary sections: a holding chamber, a flying chamber, and two attractant chambers. All elements were manufactured of Plexiglas to provide strength and transparency (Appendix A).

The holding and attractant chambers were cylindrical tubes with diameters of 4.4 cm and lengths of 12.5 cm. Each tube included a sliding door and a mesh cover at the end. The flying chamber, a square box measuring 25 × 25 × 25 cm, connected the holding chamber to the two attractant chambers by cylinder tubes (Appendix A) [21].

#### Preparation of hydrogel bead

The hydrogel beads were obtained from the Medical Entomology Laboratory, School of Biological Sciences (SBS), main campus, Universiti Sains Malaysia (USM) (Appendix B). Their preparation process is now patented (MY-195446-A). To achieve constant concentration across samples, the beads were dissolved and diluted in distilled water to 6 °Bx of the final products before being steeped in banana extract solutions for 24 hours before use (Appendix B).

#### Bioassay of Aedes aegypti

The bioassay assessing mosquito responses to attractants followed the protocol of [21,19]. We used a choice assay design with two comparison types: attractant versus control, and attractants prepared via different extraction methods. Each assay included four replicates, using 10 male and 10 female mosquitoes (5–7 days old) per replicate.

Starved mosquitoes (24 h) were acclimatized in a holding chamber for 30 min. Meanwhile, 30 g hydrogel bait loaded with fruit extract was placed in one attractant chamber, and a cotton ball soaked in 30 mL of 10% sucrose served as the control in the other. A fan ran continuously at the attractant chamber end. Sliding doors opened sequentially (attractant first for 30 s, then holding chamber), releasing mosquitoes into the flight chamber. After 4 min, the doors closed, and mosquito counts per chamber were recorded.

Chambers were washed and dried between tests to eliminate residual odours. Fresh mosquito batches were used each time, with attractant/control positions alternated across replicates to prevent side bias. Assays were repeated for all attractant pairs in Table 1.

**Table 1.**
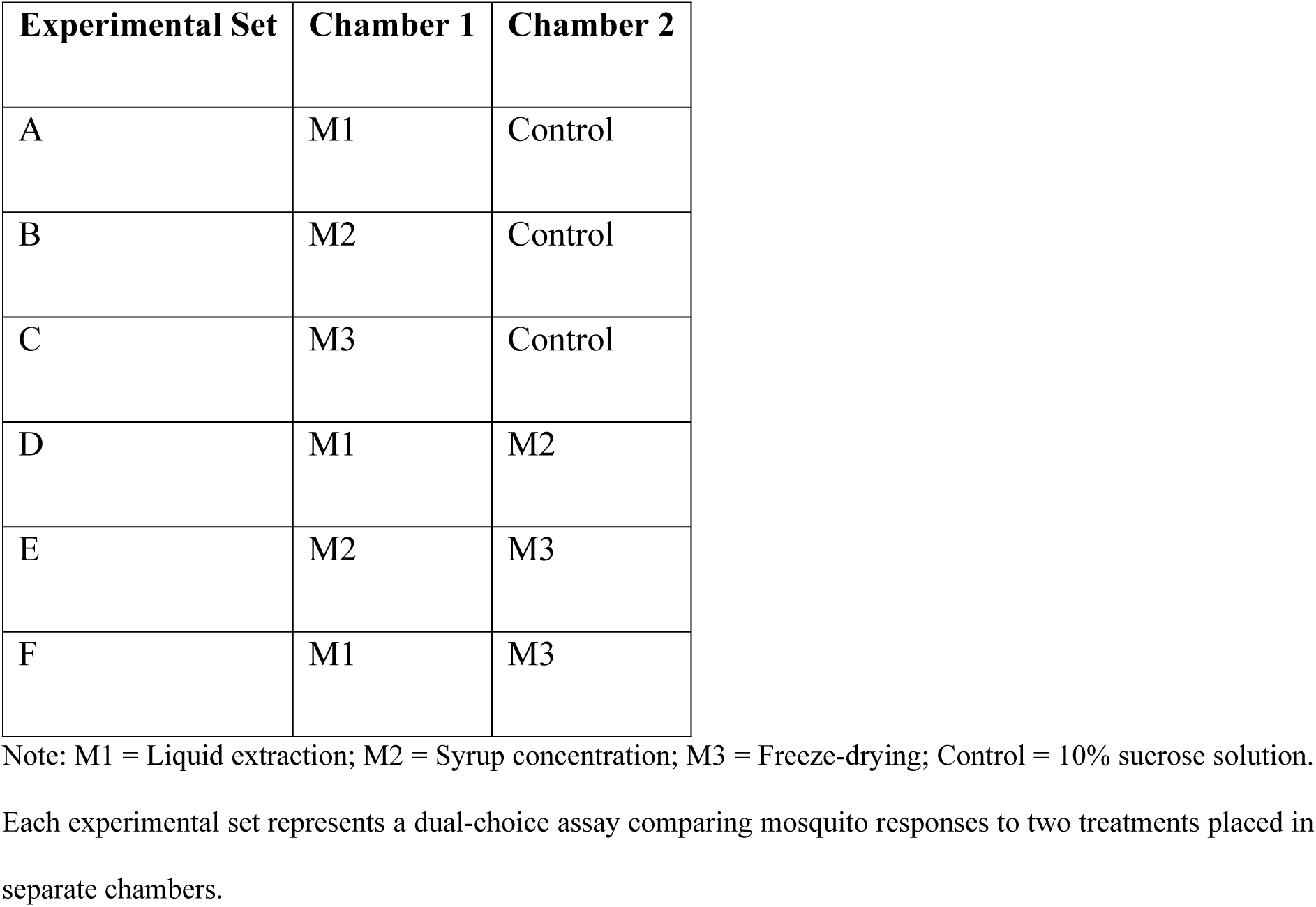
Bioassay of Aedes aegypti response toward overripe banana extract prepared using different production techniques.

| **Experimental Set** | **Chamber 1** | **Chamber 2** |
|---|---|---|
| A | M1 | Control |
| B | M2 | Control |
| C | M3 | Control |
| D | M1 | M2 |
| E | M2 | M3 |
| F | M1 | M3 |

Note: M1 = Liquid extraction; M2 = Syrup concentration; M3 = Freeze-drying; Control = 10% sucrose solution. Each experimental set represents a dual-choice assay comparing mosquito responses to two treatments placed in separate chambers.

#### Analysis of volatile constituents in banana extracts

Overripe banana extracts were analyzed for volatile chemicals by headspace gas chromatography-mass spectrometry (HS-GC/MS) using a Shimadzu® GC-2010 Plus system following the technique of [34], with minor differences.

First, banana extracts made using various manufacturing procedures were diluted with distilled water until they reached 6 °Brix (°Bx). Then, 5 mL of each diluted sample was transferred to vials and firmly sealed with PTFE-silicone septa (Appendix F)

To release volatile chemicals into the headspace, the vials were heated at 90 °C for 45 minutes and shaken continuously. The gas from the headspace was fed into the GC at 250 °C in splitless mode.

An SH-Rxi-5 Sil MS column (30 m × 0.25 mm, 0.25 μm film thickness) was employed, with helium as the carrier gas flowing at 1.5 mL per minute. The GC oven temperature was set at 50 °C, gradually raised to 300 °C at a rate of 3 °C per minute, and maintained for 5 minutes. The GC transfer line remained at 130 °C during the operation.

#### Analysis of sugar constituents in banana extracts

The quantification of sugar components in overripe banana extracts was performed at the Nutrition Laboratory, School of Health Sciences, USM Health Campus, Kelantan. The sugar components were quantified with the Boehringer Mannheim/R Biopharm sugar analysis kit for the enzymatic assessment of sucrose, glucose, fructose, and maltose. One gram of banana extract was dissolved in 60 mL of distilled water and thereafter incubated at 70 °C for 15 minutes. 5 mL of potassium hexacyanoferrate (II) solution (K₄ [Fe (CN)₆]) and 5 mL of zinc sulphate solution (ZnSO₄) were included as Carrez reagents to elucidate the sample. The pH of the mixture was modified to the range of 7.5–8.5 and subsequently precipitated for one hour. The sample solution was subsequently adjusted to a final volume of 100 mL using distilled water and centrifuged at 300 rpm for 10 minutes. The transparent supernatant was subsequently utilized for sugar analysis, adhering to the methodology provided by the sugar kit manufacturer. A UV spectrophotometer (Biochrom® WPA Biowave II) was employed to quantify the absorbance of the ensuing reaction mixtures at 340 nm, with distilled water as the blank reference. Each banana extract generated by the various manufacturing procedures was measured in triplicate.

### Statistical analysis

A Preference Index (PI) was used to access the attractiveness of overripe banana extracts prepared using different production techniques toward the *Aedes aegypti* and *Aedes albopictus.* The PI values ranged from −1 to 1, where a positive PI value indicates attraction toward the given attractants, 0 indicates neutral response, and a negative PI indicates repellence. The Preference Index (PI) was calculated using the formula:

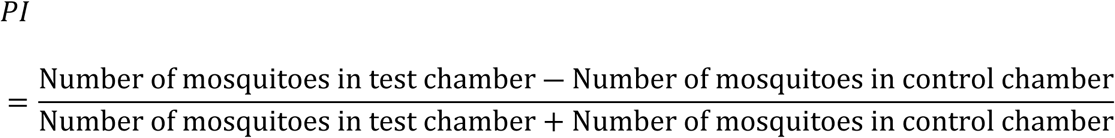

Since the Shapiro-Wilk test indicated non-normal distribution, a non-parametric Mann-Whitney U test was used to compare the attractiveness among the different attractants. Moreover, the mean of relative abundance of volatile compounds detected in overripe banana extracts using different production techniques was supported with one-way analysis of variance (ANOVA) and Tukey’s Honestly Significant Difference (HSD) test. Additionally, the Welch’s ANOVA was used to analyse the data that violated the assumption of equal variance in ANOVA. Lastly, the mean sugar content of overripe banana extracts from each production technique was also analysed using ANOVA at the significance level, α = 0.05. Once significant differences were found, Tukey’s HSD test was applied for post hoc comparisons among the treatments and the relative abundance of the volatiles. All statistical analyses were performed using IBM SPSS Statistics, Version 28.0 (IBM Corp., Armonk, NY, USA).

## Results

### Mosquito attraction bioassay Bioassay of *Aedes aegypti*

The Preference Index (PI) for *Aedes aegypti* was calculated from dual choice tests using six different combinations of attractants (Table 1). The average Preference Index (PI) values for the different banana extract types are presented in Figure 1.

**Figure 1.**
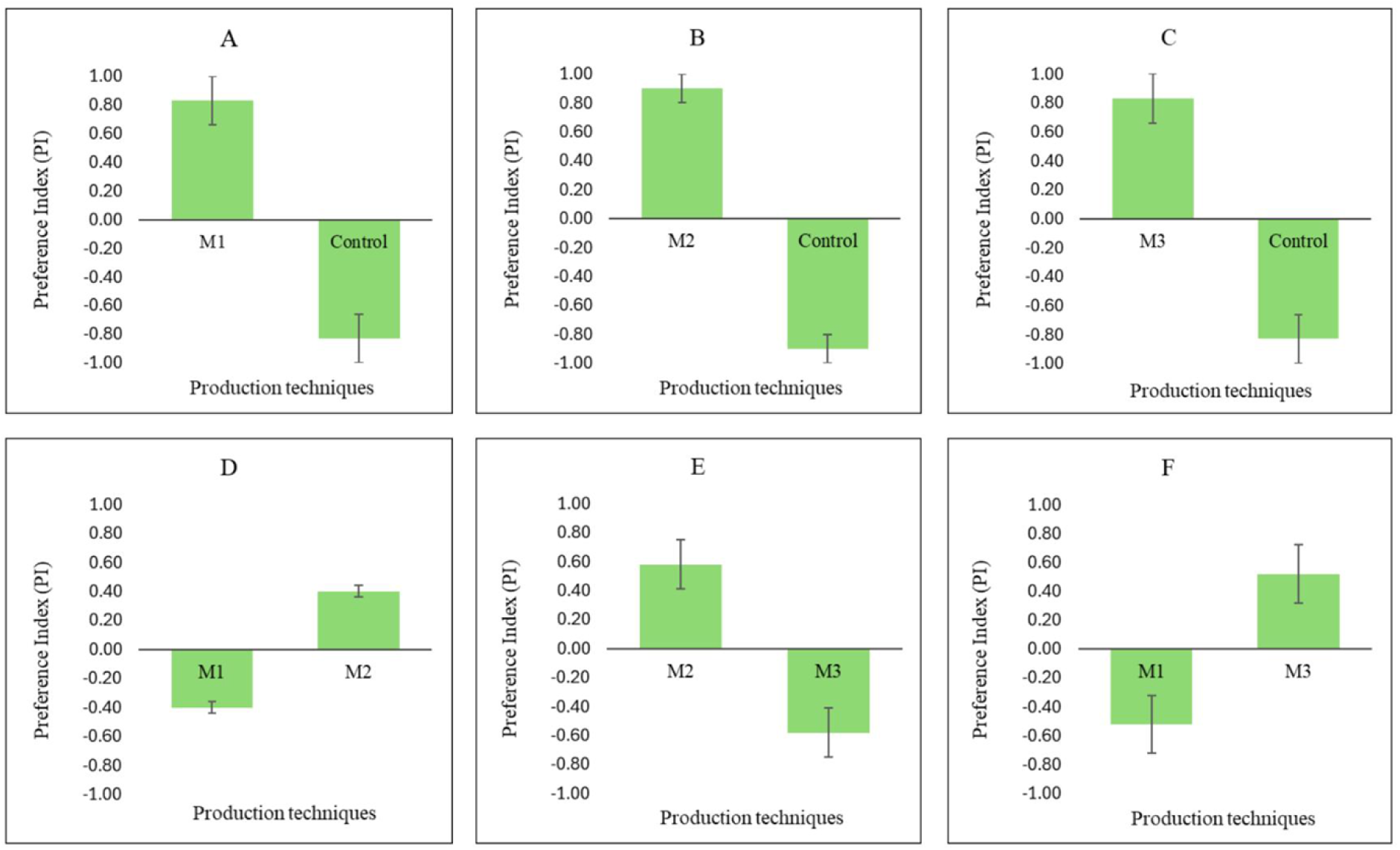
Mean Preference Index (PI ± SE) of *Aedes aegypti* in choice assay toward overripe banana extracts prepared using different production techniques, where M1: Liquid extraction; M2: Syrup concentration; M3: Freeze-drying; Control: 10% sucrose solution. (A–C) shows PI comparison between each technique and the control. (D–F) compare PI values between methods: (D) M1 vs M2, (E) M2 vs M3, and (F) M1 vs M3. The bars represent the mean ± standard error (SE).

In Set A (Figure 1(A)), *Aedes aegypti* showed a higher attraction to the banana liquid extract (PI = 0.83 ± 0.17) compared to the control. Although the data were slightly non normal, the Mann–Whitney U test still showed a significant difference (U = 0, Z = –2.530, p = 0.011).

In Set B (Figure 1(B)), the mosquitoes were strongly attracted to the banana concentrated syrup (PI = 0.90 ± 0.10), which was also significantly different from the control (U = 0, Z = –2.530, p = 0.011).

In Set C (Figure 1(C)), there was a clear and significant difference between the banana freeze dried powder (0.83 ± 0.17) and the control group (–0.83 ± 0.17) (U = 0, Z = –2.530, p = 0.011).

In Set D (Figure 1(D)), the banana concentrated syrup showed a slightly positive PI (0.40 ± 0.04), while the banana liquid extract showed a negative PI (–0.40 ± 0.04). This difference was statistically significant (U = 0, Z = –2.477, p = 0.013).

In Set E (Figure 1(E)), *Aedes aegypti* preferred the concentrated syrup (PI = 0.58 ± 0.17) over the freeze-dried powder, and this difference was significant (U = 0, Z = –2.460, p = 0.014).

In Set F (Figure1(F)), mosquitoes showed a stronger attraction to the freeze-dried powder (PI = 0.52 ± 0.20) compared to the liquid extract. This was also statistically significant (U = 0, Z = –2.477, p = 0.013).

In summary, *Aedes aegypti* showed positive attraction toward all types of overripe banana extracts when compared with the control. Among the three preparation methods, the banana concentrated syrup was the most attractive to the mosquitoes, followed by the freeze-dried extract, while the liquid extract ranked the lowest.

### Bioassay of Aedes albopictus

The average Preference Index (PI) values for *Aedes albopictus* responding to overripe banana extracts made using different production techniques are shown in Figure 2. Statistical differences for all attractant pairs were tested using the Mann–Whitney U test.

**Figure 2:**
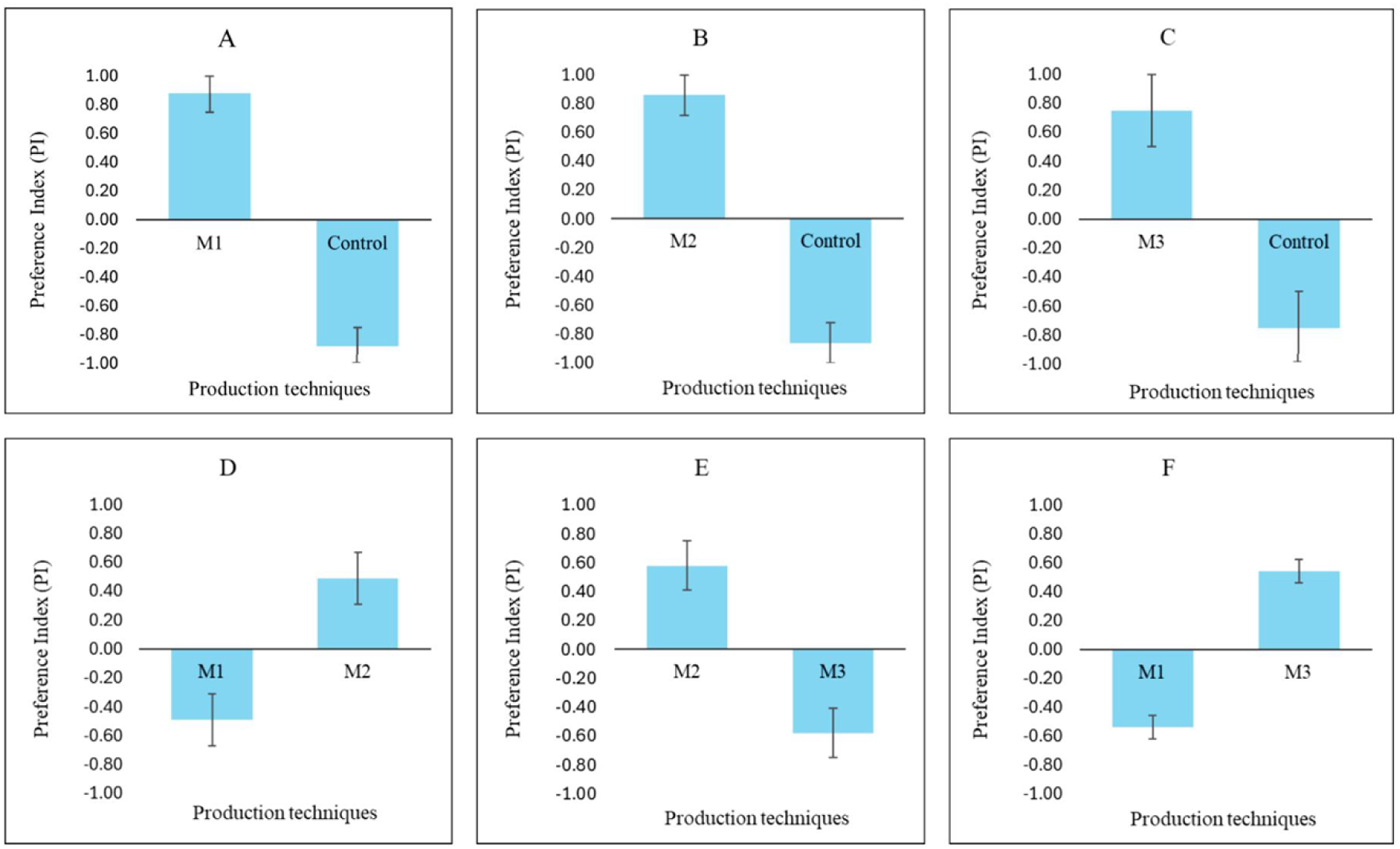
Mean Preference Index (PI ± SE) of *Aedes albopictus* in choice assay toward overripe banana extracts prepared using different production techniques, where M1: Liquid extraction; M2: Syrup concentration; M3: Freeze-drying; Control: 10% sucrose solution. (A–C) shows PI comparison between each technique and the control. (D–F) compare PI values between methods: (D) M1 vs M2, (E) M2 vs M3, and (F) M1 vs M3. The bars represent the mean ± standard error (SE).

In Set A (Figure 2 (A)), *Ae. albopictus* showed much stronger attraction to the banana liquid extract (PI = 0.88 ± 0.13) compared to the control, which had a negative PI (–0.88 ± 0.13). This difference was significant (U = 0, Z = –2.530, p = 0.011).

In Set B (Figure 2 (B)), the mosquitoes also showed a significant preference for the banana concentrated syrup (PI = 0.86 ± 0.14) compared to the control group (U = 0, Z = –2.530, p = 0.011).

In Set C (Figure 2 (C)), attraction to the banana freeze dried powder (0.75 ± 0.25) was significantly higher than the control (–0.75 ± 0.25) (U = 2.000, Z = –2.049, p = 0.040).

In Set D (Figure 2 (D)), *Ae. albopictus* preferred the banana concentrated syrup (PI = 0.49 ± 0.18) over the banana liquid extract. This difference was significant (U = 0, Z = –2.460, p = 0.014).

In Set E (Figure 2 (E)), the banana concentrated syrup again showed a higher attraction value (0.58 ± 0.14) compared to the freeze-dried extract (U = 0, Z = –2.477, p = 0.013).

In Set F (Figure 2 (F)), mosquitoes were significantly more attracted to the banana freeze dried powder (PI = 0.54 ± 0.08) than to the banana liquid extract, which had a negative PI (–0.54 ± 0.08) (U = 0, Z = –2.477, p = 0.013).

In summary, just like *Aedes aegypti*, *Aedes albopictus* was more attracted to all types of overripe banana extracts compared to the control. When comparing the extracts to each other, the banana concentrated syrup consistently showed the strongest attraction, followed by the freeze-dried extract, with the liquid extract showing the lowest attraction. These findings support the idea that the way banana extracts are produced plays an important role in how attractive they are to mosquitoes.

### Volatile Constituents in Banana Extracts

In total, 16 volatile organic compounds (VOCs) were found in the overripe banana extracts produced using the different techniques (Table 2). These compounds match what has been reported in earlier studies on banana volatiles [35,36, 37,8, 19]. All VOCs were grouped into five categories: esters, alcohols, aldehydes, ketones, and others. Among these, esters were the most dominant, especially in the freeze-dried powder (M3). A one-way ANOVA was used to test the significant differences between techniques.

**Table 2:**
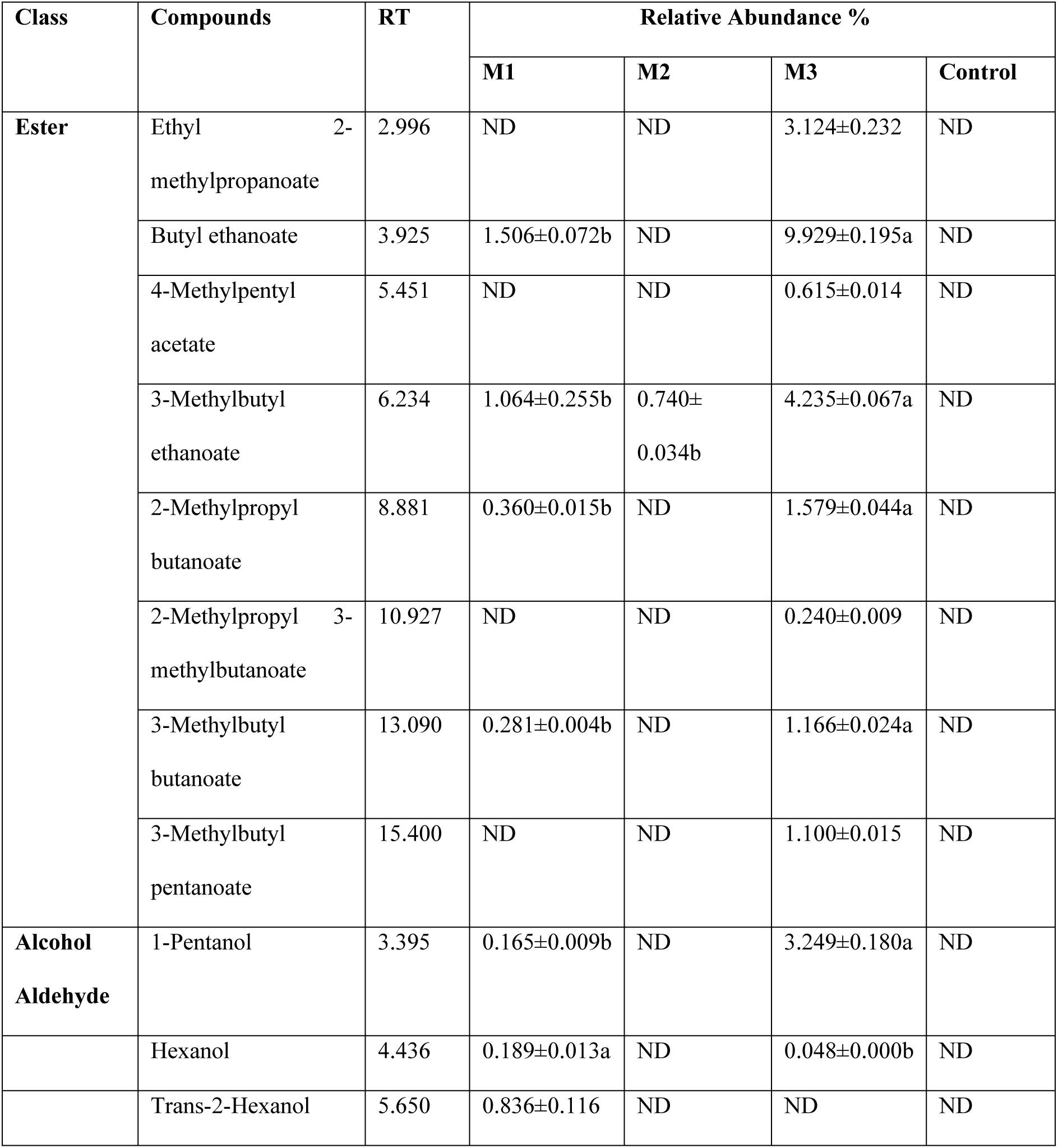
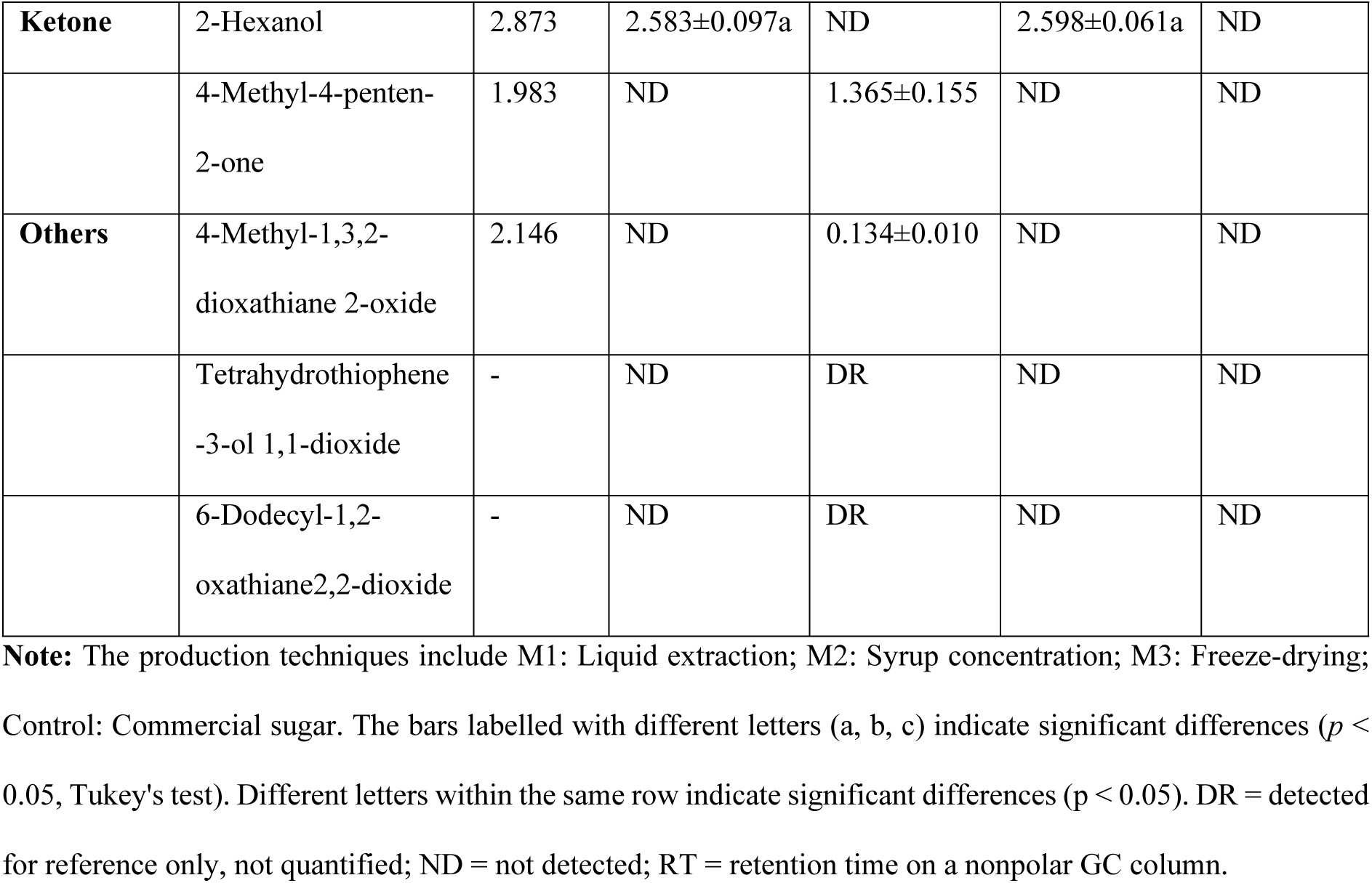
Relative abundances of volatile organic compounds (VOCs) in overripe banana extract prepared using different production techniques.

| Class | Compounds | RT | Relative Abundance % |  |  |  |
|---|---|---|---|---|---|---|
|  |  |  | M1 | M2 | M3 | Control |
| **Ester** | Ethyl 2-methylpropanoate | 2.996 | ND | ND | 3.124±0.232 | ND |
|  | Butyl ethanoate | 3.925 | 1.506±0.072b | ND | 9.929±0.195a | ND |
|  | 4-Methylpentyl acetate | 5.451 | ND | ND | 0.615±0.014 | ND |
|  | 3-Methylbutyl ethanoate | 6.234 | 1.064±0.255b | 0.740±0.034b | 4.235±0.067a | ND |
|  | 2-Methylpropyl butanoate | 8.881 | 0.360±0.015b | ND | 1.579±0.044a | ND |
|  | 2-Methylpropyl 3-methylbutanoate | 10.927 | ND | ND | 0.240±0.009 | ND |
|  | 3-Methylbutyl butanoate | 13.090 | 0.281±0.004b | ND | 1.166±0.024a | ND |
|  | 3-Methylbutyl pentanoate | 15.400 | ND | ND | 1.100±0.015 | ND |
| **Alcohol Aldehyde** | 1-Pentanol | 3.395 | 0.165±0.009b | ND | 3.249±0.180a | ND |
|  | Hexanol | 4.436 | 0.189±0.013a | ND | 0.048±0.000b | ND |
|  | Trans-2-Hexanol | 5.650 | 0.836±0.116 | ND | ND | ND |
| **Ketone** | 2-Hexanol | 2.873 | 2.583±0.097a | ND | 2.598±0.061a | ND |
|  | 4-Methyl-4-penten-2-one | 1.983 | ND | 1.365±0.155 | ND | ND |
| **Others** | 4-Methyl-1,3,2-dioxathiane 2-oxide | 2.146 | ND | 0.134±0.010 | ND | ND |
|  | Tetrahydrothiophene-3-ol 1,1-dioxide | - | ND | DR | ND | ND |
|  | 6-Dodecyl-1,2-oxathiane2,2-dioxide | - | ND | DR | ND | ND |

**Note:** The production techniques include M1: Liquid extraction; M2: Syrup concentration; M3: Freeze-drying; Control: Commercial sugar. The bars labelled with different letters (a, b, c) indicate significant differences (*p* < 0.05, Tukey's test). Different letters within the same row indicate significant differences (p < 0.05). DR = detected for reference only, not quantified; ND = not detected; RT = retention time on a nonpolar GC column.

When looking at the number of compounds, the freeze-dried powder had the highest diversity with 11 VOCs, followed by the liquid extract (M1) with 8 VOCs, and the concentrated syrup (M2) with 5 VOCs. The control sample showed no detectable volatile compounds.

Among the ester compounds, 3 methylbutyl ethanoate (isoamyl acetate) was found in all three banana extract types but not in the control. A one-way ANOVA showed a significant difference between techniques for this ester (F (2,6) = 385.613, p < 0.001). Tukey’s HSD test showed that the freeze-dried powder had a much higher level of this compound (4.235 ± 0.067%) than both the liquid extract and concentrated syrup (p < 0.001).

Another ester, butyl ethanoate, was the most abundant compound overall. The freeze-dried extract showed a significantly higher amount (9.929 ± 0.195%) compared to the liquid extract (1.506 ± 0.072%). Similarly, 2 methylpropyl butanoate and 3 methylbutyl butanoate were both found at much higher levels in the freeze-dried sample than in the liquid extract. Some esters for example ethyl 2 methylpropanoate, 4 methylpentyl acetate, and 3 methylbutyl pentanoate were found only in the freeze-dried powder.

For the alcohol group, only 1 pentanol was detected. The freeze-dried powder had a much higher amount (3.429 ± 0.180%) than the liquid extract (0.165 ± 0.009%). This difference was significant according to Welch’s ANOVA (F (1,2.004) = 189.61, p = 0.005).

For aldehydes, hexanal was more abundant in the liquid extract (0.189 ± 0.013%) compared to the freeze-dried powder (0.048 ± 0.000%). For ketones, 2 hexanone appeared in both the liquid extract (2.583 ± 0.097%) and freeze-dried powder (2.598 ± 0.061%), with no significant difference between them (ANOVA, F (1,4) = 0.012, p = 0.919).

The concentrated syrup had fewer VOCs, most likely because some compounds were lost during heating. However, it contained new thermally formed compounds such as 4 Methyl 4 penten 2 one and 4 methyl 1,3,2 dioxathiane 2 oxide which were not detected in the other extracts. These compounds may have been produced by Maillard reactions during the heating process.

Overall, the differences in VOC profiles agreed with the mosquito bioassay results, showing that both the amount and type of volatile compounds can influence mosquito attraction.

### Sugar constituents in banana extracts

The mean sugar contents (± SE) of sucrose, glucose, fructose, and maltose from each production technique are shown in Figures 3 to 6. All sugar data were analysed using one way ANOVA.

**Figure 3:**
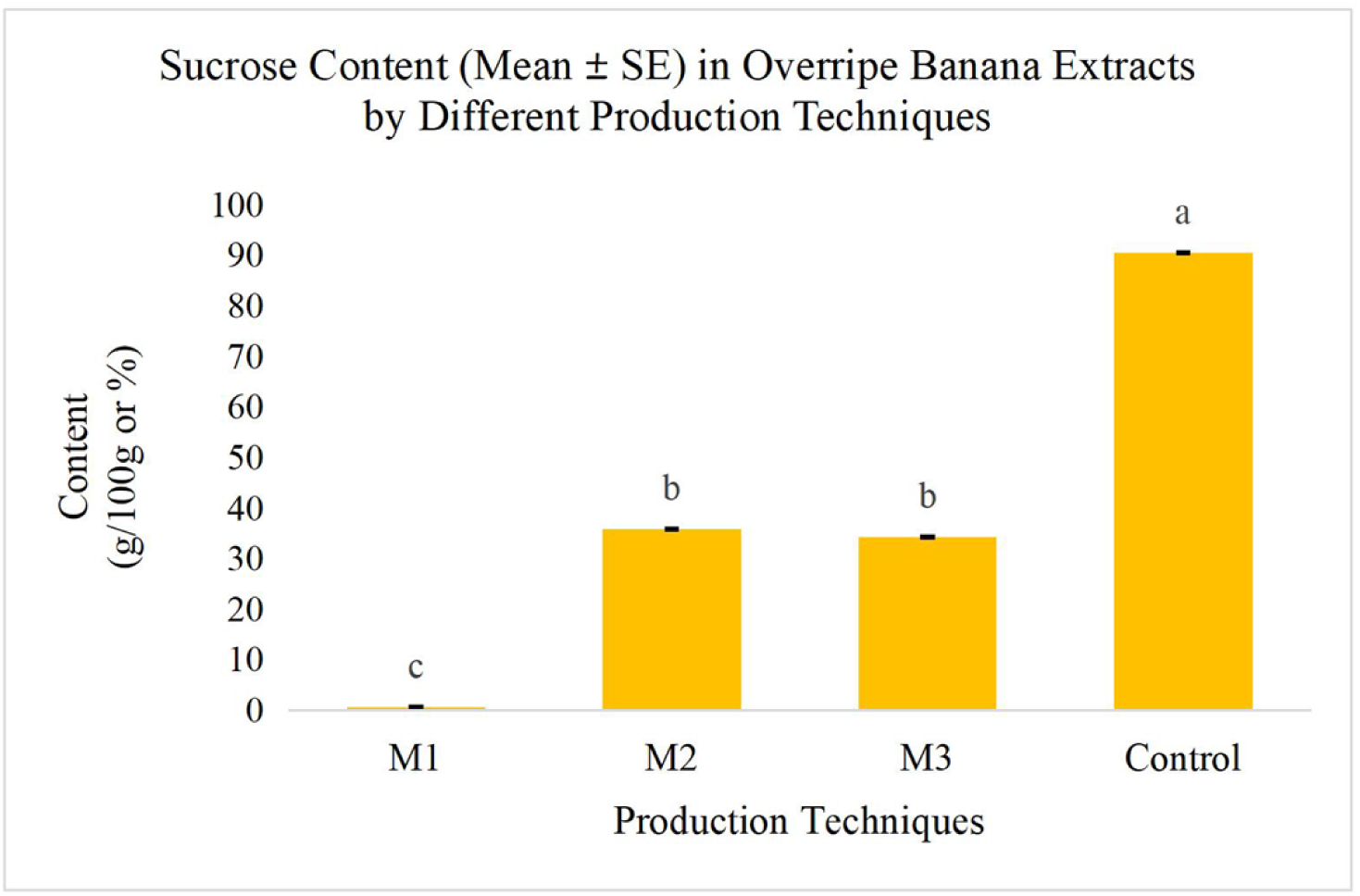
Mean ± SE of sucrose content in overripe banana extracts prepared using different production techniques, where M1: Liquid extraction; M2: Syrup concentration. M3: Freeze-drying; Control: Commercial sugar. The bars labelled with different letters (a, b, c) indicate significant differences (*p* < 0.05, Tukey’s test).

For sucrose, Welch’s one way ANOVA showed a significant difference among the different production techniques (F (3, 3.49) = 7136.35, p < 0.001). The Games–Howell post hoc test confirmed that all groups were significantly different from one another (p < 0.05), except for the concentrated syrup and freeze-dried powder, which were not significantly different (p =0.885).

As shown in Figure 3, the sucrose content of the liquid extract was 0.57 ± 0.12 g/100 g, while the concentrated syrup contained 35.76 ± 1.08 g/100 g, and the freeze-dried powder contained 34.20 ± 1.90 g/100 g. The sugar control had the highest sucrose level at 90.43 ± 0.52 g/100 g.

Overall, the commercial sugar control showed the highest sucrose content, whereas the liquid extraction method produced the lowest sucrose level.

For glucose content, a one-way ANOVA showed that there was a significant difference among the different production techniques (F (3, 8) = 1393.07, p < 0.001). As shown in Figure 4, the freeze-dried powder had the highest glucose level at 28.70 ± 0.44 g/100 g, and this value was significantly different from all other groups according to Tukey’s HSD test (p < 0.05). On the other hand, the banana liquid extract had the lowest glucose content at 2.16 ± 0.37 g/100 g, and it was not significantly different from the commercial sugar control, which had 3.29 ± 0.10 g/100 g (p = 0.187). The concentrated syrup recorded 21.20 ± 0.40 g/100 g, which was significantly lower than the freeze-dried powder but still much higher than the liquid extract.

**Figure 4.**
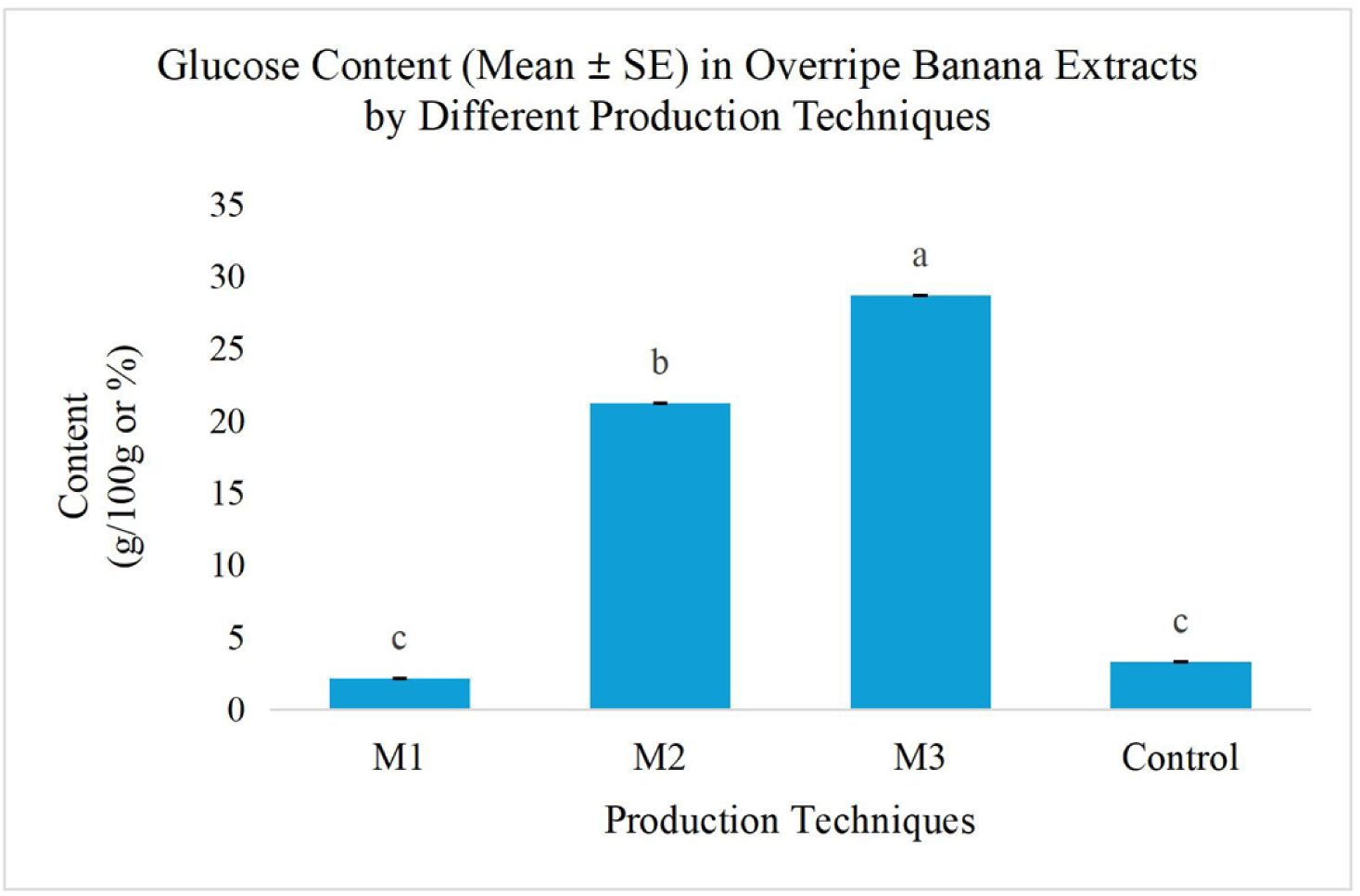
Mean of glucose content in overripe banana extracts prepared using different production techniques, where M1: Liquid extraction; M2: Syrup concentration; M3: Freeze-drying; Control: Commercial sugar. The bars labelled with different letters (a, b, c) indicate significant differences (*p* < 0.05, Tukey’s test).

**Figure 5:**
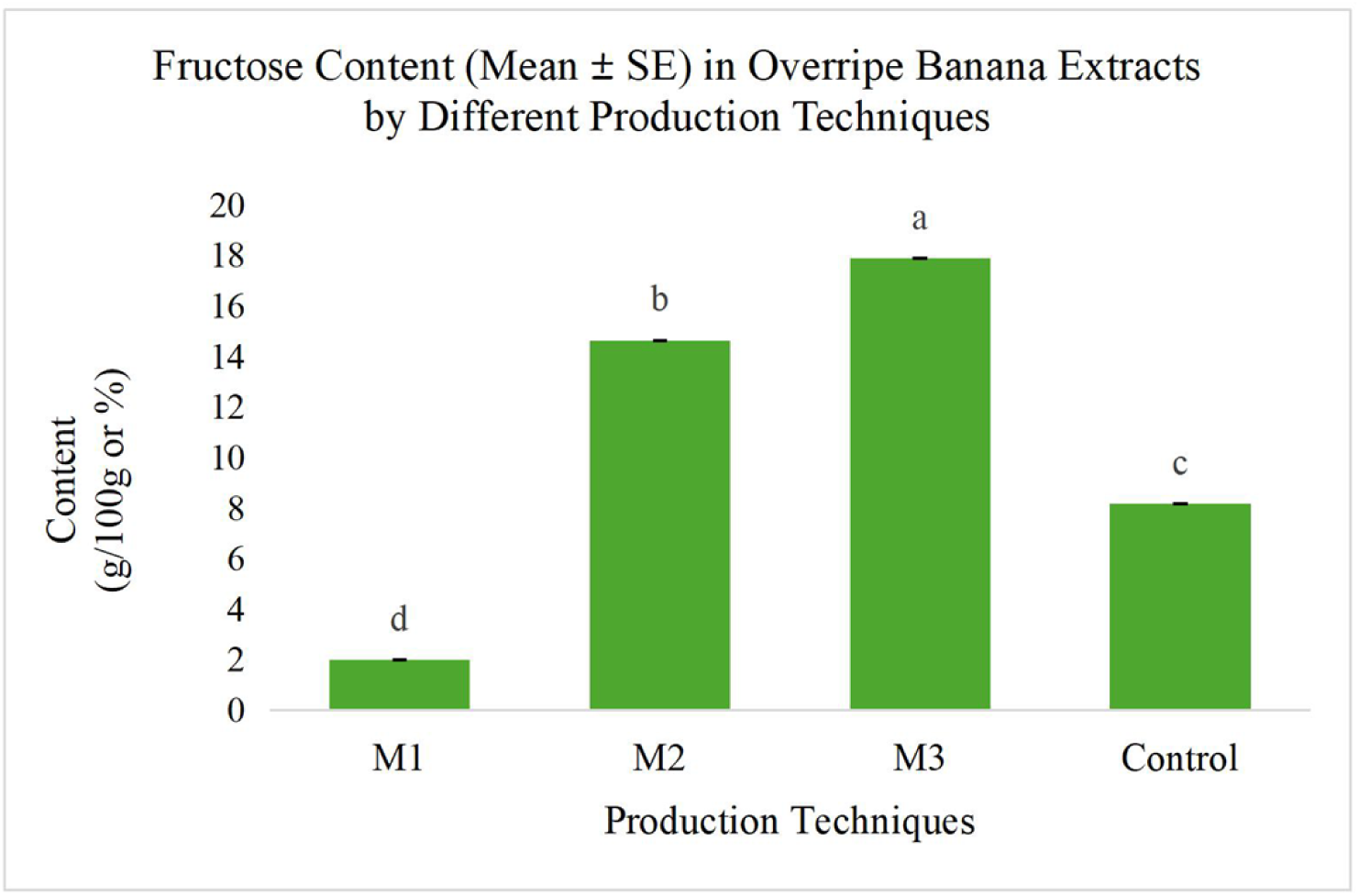
Mean ± SE of fructose content in overripe banana extracts prepared using different production techniques, where M1: Liquid extraction; M2: Syrup concentration. M3: Freeze-drying; Control: Commercial sugar. The bars labelled with different letters (a, b, c, d) indicate significant differences (*p* < 0.05, Tukey’s test).

For maltose content, there was a statistically significant difference within the production techniques (Welch’s ANOVA, F (3,4.06) = 131.54, p < 0.001). According to Figure 6, freeze-drying recorded the highest maltose content (15.56 ± 1.45 g/100g) but was not significantly different from the syrup concentration (9.28 ± 0.70 g/100g) (p = 0.09). In contrast, the commercial sugar resulted in the lowest (−2.97 ± 0.18 g/100g), which was significantly different from all other groups based on the Games-Howell test (p < 0.05). The liquid extract (0.59 ± 0.19 g/100g) also recorded a relatively low level of maltose.

The variations in sugar content observed among the different banana extracts correspond with the mosquito preference patterns recorded at the bioassays. The high levels of natural sugars, particularly glucose and fructose, were associated with increased mosquito attraction.

In conclusion, the results demonstrated that the production technique had a significant impact on the chemical composition of overripe banana extracts. The syrup concentration method notably enhances sugar levels and may lead to the formation of additional volatile compounds, while the freeze-drying method effectively preserves a wide range of volatile organic compounds (VOCs) and natural sugars. These chemical properties were consistent with the observed increase in mosquito attraction.

## Discussion

The present study demonstrated that processing techniques significantly influenced the attractiveness of overripe banana (*Musa acuminata*) extracts to *Aedes aegypti* and *Aedes albopictus*. Both mosquito species exhibited significantly greater attraction to banana-derived formulations than to the 10% sucrose control. Among the three preparation methods, concentrated syrup consistently produced the strongest attraction, followed by freeze-dried and liquid extracts. These findings highlight the importance of processing-dependent changes in fruit composition in determining mosquito olfactory preferences. Mosquitoes rely on plant-derived volatile organic compounds (VOCs) to locate potential sugar sources, and recent investigations have identified several odorant receptors in *Ae. aegypti* that respond to floral and plant-associated volatiles [38]. The greater attractiveness of banana extracts compared with sucrose alone may therefore be associated with the presence of complex fruit-derived odour blends that provide olfactory cues for mosquito orientation.

Previous studies have demonstrated that mosquito attraction varies considerably among fruit species and their volatile profiles. Leon-Noreña et al [26] reported that, *Ae. aegypti* exhibited selective preferences for different fruit odours, including banana, mango, peach, and guava, and demonstrated that changes in the relative proportions of volatile constituents could modify attractiveness. Similarly, Ayub et al [19] investigated fruit-based attractants for hydrogel bait applications targeting *Aedes* mosquitoes, highlighting the potential of fruit combinations in attractive sugar bait formulations. These observations support the present findings and suggest that the attractiveness of banana extracts is associated not merely with the presence of fruit volatiles but also with the composition and relative proportions of compounds within their odour blends.

Interestingly, concentrated syrup exhibited the strongest attraction in both mosquito species despite containing fewer detectable VOCs than the freeze-dried extract. This finding suggests that mosquito attraction may depend more strongly on the identity, concentration, relative proportions, and interactions of behaviourally active compounds than on overall VOC diversity. Hutcheson and his research team demonstrated that, *Ae. aegypti* and *Ae. albopictus* responded differently to synthetic phytochemical blends depending on their component ratios and concentrations [39]. Similarly, Leon-Noreña et al [26] demonstrated that relatively small changes in fruit odour composition could alter mosquito attractiveness. Together, these studies provide a plausible explanation for the stronger attraction to syrup-concentrated banana extracts despite their comparatively limited detected volatile profile. Thermal concentration may have modified the composition and release characteristics of the banana odour blend through the loss, transformation, or formation of particular compounds. Although some compounds were detected exclusively in the concentrated syrup, their formation pathways and individual contributions to mosquito attraction remain to be established.

Freeze-dried extracts exhibited the second-highest attractiveness and contained the most diverse VOC profile, particularly esters associated with characteristic banana aromas. Among the detected compounds, butyl ethanoate (9.929%) and 3-methylbutyl ethanoate (isoamyl acetate; 4.235%) were substantially more abundant in the freeze-dried extract than in the liquid extract. Additional esters, including ethyl 2-methylpropanoate, 4-methylpentyl acetate, and 3-methylbutyl pentanoate, were detected exclusively in the freeze-dried preparation. These observations are consistent with [40], who investigated the volatile composition of banana powders produced using different drying techniques and identified esters as major contributors to characteristic banana aromas. The retention of a relatively diverse ester profile following freeze-drying may therefore have contributed to the greater attractiveness of this formulation compared with the liquid extract. However, the lower attraction of freeze-dried extract relative to concentrated syrup indicates that greater detected VOC diversity does not necessarily correspond to stronger mosquito attraction.

Among the identified compounds, isoamyl acetate is of particular interest because of its characteristic banana-like aroma and its potential role in mosquito responses to fruit odours. Leon-Noreña et al [26] demonstrated that, *Ae. aegypti* attraction to fruit odour blends can be altered by changes in the proportions of individual volatile constituents, supporting the importance of specific compounds within complex mixtures. Other esters identified in the present study, including butyl ethanoate, 2-methylpropyl butanoate, and 3-methylbutyl butanoate, may also contribute to the overall olfactory characteristics of banana extracts. Nevertheless, the detection of these compounds does not establish their individual attractiveness to mosquitoes. Their behavioural significance may depend on their concentrations, interactions with other volatile constituents, and release rates from the hydrogel matrix. Further investigation using individual compounds and reconstructed synthetic blends would be necessary to determine their specific contributions to mosquito attraction.

The liquid extract consistently exhibited the lowest attractiveness among the three banana preparations. This response may be associated with its comparatively limited ester profile and lower relative abundances of several fruit-associated volatile constituents. Previous research has demonstrated that drying and thermal processing can substantially alter banana aroma composition. Saha et al [34] reported that low-temperature drying resulted in considerable losses of banana flavour volatiles, particularly esters, with reductions varying from 25% to 87% under the investigated conditions. Similarly, Wang et al demonstrated that drying techniques influence the composition of banana aroma compounds [40]. These findings suggest that differences in volatile retention during processing may contribute to the chemical variation observed among the present formulations. However, because the current study did not directly quantify volatile losses during each processing stage, the lower attractiveness of the liquid extract cannot be attributed conclusively to thermal degradation alone. Differences in extraction efficiency, dilution, and volatile-release characteristics may also have influenced mosquito responses.

Sugar profiling revealed substantial differences in carbohydrate composition among the banana preparations. Syrup-concentrated and freeze-dried extracts contained higher concentrations of several measured sugars than the liquid extract, whereas the commercial sugar control contained the highest sucrose concentration. Despite its high sucrose content, the control was significantly less attractive than the banana formulations. This observation suggests that sucrose concentration alone was insufficient to explain the olfactory preferences recorded in the present study. Although sugar feeding is essential for mosquito energy acquisition, survival, and flight activity, sugars such as glucose, fructose, sucrose, and maltose are essentially nonvolatile under the experimental conditions and are therefore unlikely to function directly as long-range olfactory attractants. Nevertheless, carbohydrate composition may influence formulation properties, volatile release, and subsequent feeding responses following contact with the attractant. Further investigations are needed to distinguish the contribution of olfactory cues from the nutritional characteristics of banana-based formulations.

The incorporation of banana extracts into hydrogel beads represents a potentially useful approach for developing fruit-based mosquito attractants. However, several limitations should be considered when interpreting the present findings. First, the GC–MS analysis characterised the relative abundance of volatile compounds in the extracts rather than directly measuring their emission from hydrogel beads during behavioural assays. Consequently, the detected chemical profiles may not fully represent the volatile mixtures encountered by mosquitoes within the olfactometer. Second, individual compounds were not evaluated independently, preventing identification of the specific constituents responsible for attraction. Furthermore, variations in banana ripeness, sample preparation, and hydrogel stability may influence the reproducibility of fruit-based formulations. Future investigations should therefore evaluate the headspace emissions of formulated hydrogel beads, determine the behavioural activity of selected compounds and synthetic blends, and assess formulation stability under different storage conditions.

Overall, the present findings demonstrate that banana-processing techniques significantly influence the chemical characteristics and olfactory attractiveness of overripe banana extracts to *Ae. aegypti* and *Ae. albopictus*. The greater attractiveness of concentrated syrup despite its lower detected VOC diversity suggests that mosquito responses may be determined by specific volatile constituents and their interactions rather than by overall compound diversity alone. These findings provide a scientific basis for further optimisation of banana-derived hydrogel attractants and their potential integration into attractive toxic sugar bait strategies. Further behavioural, chemical, and field-based investigations will be essential to identify the active components and evaluate their practical effectiveness in mosquito surveillance and control.

## Conclusion

This study demonstrated that different processing techniques applied to overripe banana (*Musa acuminata*) significantly influenced the attractiveness of banana extracts to *Aedes aegypti* and *Aedes albopictus*, as well as their volatile organic compound (VOC) profiles and sugar composition. Both mosquito species exhibited the strongest attraction to syrup-concentrated extracts, followed by freeze-dried and liquid extracts. GC–MS analysis revealed that freeze-drying retained the most diverse VOC profile, particularly fruit-associated esters such as isoamyl acetate and 2-methylpropyl butanoate. In contrast, syrup concentration produced fewer detectable VOCs but exhibited the highest mosquito attractiveness, suggesting that specific volatile constituents, their relative proportions, or release characteristics may be more important than overall VOC diversity. Sugar analysis further revealed higher concentrations of several carbohydrates in the syrup-concentrated and freeze-dried preparations than in the liquid extract, although their direct contribution to olfactory attraction remains uncertain.

Several practical limitations were encountered, including variability in banana ripening, susceptibility of fruit-based hydrogel beads to spoilage during storage, and difficulties in coordinating sample preparation for GC–MS analysis across the three processing methods. These factors highlight the importance of standardising raw material maturity, sample preparation, and storage conditions to improve experimental reproducibility.

Future investigations should focus on identifying and evaluating the behaviourally active VOCs responsible for the enhanced attractiveness of syrup-concentrated extracts, particularly compounds potentially associated with thermal processing. Further optimisation of banana extract concentrations, hydrogel stability, and preservation methods is also warranted. Additionally, headspace analysis of formulated hydrogel beads and validation under semi-field and field conditions would help establish their practical effectiveness. Overall, these findings highlight the potential of processed overripe banana extracts as natural attractant components for the development of hydrogel-based mosquito surveillance and control strategies.

## Authors contributions

SAA and NFAK conceived and designed the study. SAA, LWH, SS and NAAL carried out the investigation and data curation. Formal analysis was performed by SAA, LWH and NYV. NFAK and CEW secured funding for the project. Methodology was developed by SAA, LWH, NAAL and NFAK. Project administration was managed by NFAK and WRWI, while resources were provided by WRWI, CEW and NFAK. Software support was contributed by NYV and SAA. Supervision was conducted by NFAK and CEW. Validation was performed by SAA, LWH and NYV. Visualization was carried out by SAA and LWH. SAA prepared the original draft, and all authors (SAA, LWH, SS, NAAL, WRWI, NYV, CEW and NFAK) contributed to review and editing of the final manuscript.

## Acknowledgments

This research is supported by BioG Expert Sdn. Bhd. and Ministry of Higher Education Malaysia.

